# Chromosome-level genome assembly of a doubled haploid brook trout (*Salvelinus fontinalis*)

**DOI:** 10.1101/2024.09.27.615458

**Authors:** Laurie Lecomte, Anne-Laure Ferchaud, Eric Normandeau, Claire Mérot, Isabelle Langlois-Parisé, Jean-Christophe Therrien, Pierre Bérubé, Haig Djambazian, Pubudu Manoj Nawarathna, Dylan Fraser, Jiannis Ragoussis, Louis Bernatchez

## Abstract

Brook trout (*Salvelinus fontinalis*) is a socioeconomically important fish species for fisheries, aquaculture and aquatic conservation. We produced a 2.5 Gb reference assembly by combining Hi-C chromosome conformation capture with high-coverage short- and long-read sequencing of a fully homozygous mitotic gynogenic doubled haploid fish, which facilitates assembly of highly complex salmonid genomes. The assembly has a N50 of 50.98 Mb and 88.9% of the total assembled sequence length is anchored into 42 main chromosomes, of which 63.44% represents repeated contents, including 1,461,010 DNA transposons. 56,058 genes were found with the NCBI annotation pipeline, with 99% of the 3,640 expected conserved orthologs BUSCO genes (actinopterygii_odb10 lineage database). Additionally, we found significant homology within the 42 chromosomes, as expected for this pseudo-tetraploid species, as well as with the sister species lake trout (*Salvelinus namaycush*) and Atlantic salmon (*Salmo salar*). This assembly will serve as a reliable genomic resource for brook trout, thus enabling a wider range of reference-based applications to support ongoing research and management decision-making for the species.

## Introduction

Brook trout (*Salvelinus fontinalis*) is a freshwater salmonid fish native to Eastern North America (Scott & Crossman, 1973; McCrimmon & Campbell, 1969; Behnke, 2002). Here, the species supports recreational fisheries, subsistence fisheries and aquaculture, and is an indicator species for assessing cold water aquatic ecosystem health (Berkes,1979; JBNQNHRC, 1982; Ficke, Peterson & Janowsky, 2009). It has also been introduced for recreational purposes in various freshwater habitats beyond its native range in Canada and the United States, as well as in other continents (McCrimmon & Campbell, 1969; Kitano, 2004).

The evolutionary history of brook trout was shaped by glacial events leading to postglacial colonization, as inferred from mitochondrial DNA (Dantzmann et al., 1998; Pilgrim et al., 2012). At least, two phylogenetically distinct major lineages have been documented across the species’ native range. The northern strain is found in Canada and the northeastern United States. The southern strain is found in the southern Appalachian Mountains (Stoneking, Wagner & Hildebrand, 1981; McCracken, Parker & Guffey, 1993), and its populations exhibit greater interpopulation haplotype diversity and heterogeneity (Danzmann et al., 1998; Guffrey et al., 1999; Habera & Moore, 2005). Brook trout populations usually show high levels of genetic structure, often at small geographic scales, that is primarily determined by watershed (Angers & Bernatchez, 1998; Danzmann & Ihssen, 1995; Castric, Bonney & Bernatchez, 2001; Ferchaud et al., 2020; Lehnert et al., 2020) and landscape features (Castric, Bonney & Bernatchez, 2001; Wood et al., 2014; Kazyak et al., 2022).

In the last century, brook trout populations have experienced decline across their habitat range as a result of anthropogenic influences, mainly habitat destruction and alteration, climate change, exotic invasive species and overexploitation (McCrimmon & Campbell, 1969; Larson & Moore, 1985; Marschall & Crowder, 1996; Hudy et al. 2008; Lehnert et al., 2020). This is especially true for small-sized southern populations isolated in high-elevation small habitats (Kazyak et al., 2022), in which reduced gene flow and genetic diversity may negatively impact adaptive potential, making these populations more vulnerable to collapse (Keller & Waller, 2002; King et al., 2012, Ferchaud et al., 2020) and less resilient to rapid environmental change (Mathur et al., 2023).

To mitigate such decline, as well as to support recreational fishing, wild brook trout populations have been extensively supplemented with hatchery-reared individuals (McCrimmon & Campbell, 1969; Kerr, 2000), a widespread practice with highly variable and often deleterious impacts in salmonids. Indeed, supplementation with hatchery-reared fish derived from nonlocal broodstock may lead to introgression, which may threaten population-specific adaptive variation (Allendorf et al., 2001; Laikre et al., 2010). Hatchery-origin fish also reportedly have lower fitness in the wild than native local individuals (Araki et al., 2008; Christie et al., 2014). In addition to these traditional management tools, genetic rescue has been recently pointed out as one of the only approaches that can provide variation needed for future adaptation to changing conditions and that could thus reduce the rate of contemporary brook trout extirpation (White et al., 2023). Consequently, knowledge of phylogeographic relationships and population structure is crucial for a variety of conservation initiatives for brook trout, including population management and selection of adequate hatchery broodstocks.

As a part of the Canada 150 Sequencing Initiative (CanSeq150), we generated a high-quality reference genome assembly by combining high coverage short- and long-read sequencing with Hi-C chromosome conformation capture. To maximize assembly contiguity and precision, we used a homozygous, doubled haploid fish we produced through mitotic gynogenesis (Zhang et al., 2014; Biagi et al., 2022). This technique offered an ideal context to aid the assembly of such a complex salmonid genome with a size of 2.5 Gb, a high proportion of repetitive contents (50 to 60%; de Boer et al., 2007), as well as numerous duplicated regions and residual tetrasomy resulting from the whole genome duplication event that occurred at least 60 million years ago in salmonids (Allendorf & Thorgaard, 1984; Crête-Lafrenière, Weir & Bernatchez, 2012; Lien et al., 2016). Experimental details are presented here to provide a methodological resource for the production of doubled haploid samples in other species with complex genomes. This assembly represents a valuable resource that will allow whole-genome-scale characterization of polymorphism, thus revealing a previously unexplored layer of genetic variation and yielding new insights on brook trout adaptation and evolution.

## Methods

### Doubled haploid production

Doubled haploids are produced by retaining duplicated set of chromosomes from a parental haploid cell, yielding fully homozygous individuals with genetic material inherited from a single parent (Jagiełło et al., 2021). To produce mitotic gynogens, i.e., doubled haploids with two identical copies of each maternal chromosome, haploid eggs are fertilized with sperm carrying deactivated DNA, then exposed to thermal or pressure shock to disturb the first cellular division and retain the duplicated maternal chromosome pair (Figure 1a) (Biagi et al., 2022). Doubled haploid brook trout were reared at Université Laval’s Laboratoire de recherche en sciences environnementales et médicales (LARSEM). Given that optimal experimental parameters for mitotic gynogenesis have yet to be determined for brook trout, three UV irradiation doses and five pressure shock times were trialled. Each egg group was subjected to a different combination of a given UV irradiation dose and a given pressure shock time, yielding 19 treatment and control groups (Figure 1b). The procedure was authorized by the Comité de protection des animaux de l’Université Laval (permit number: 2018111-1).

**Figure 1.**
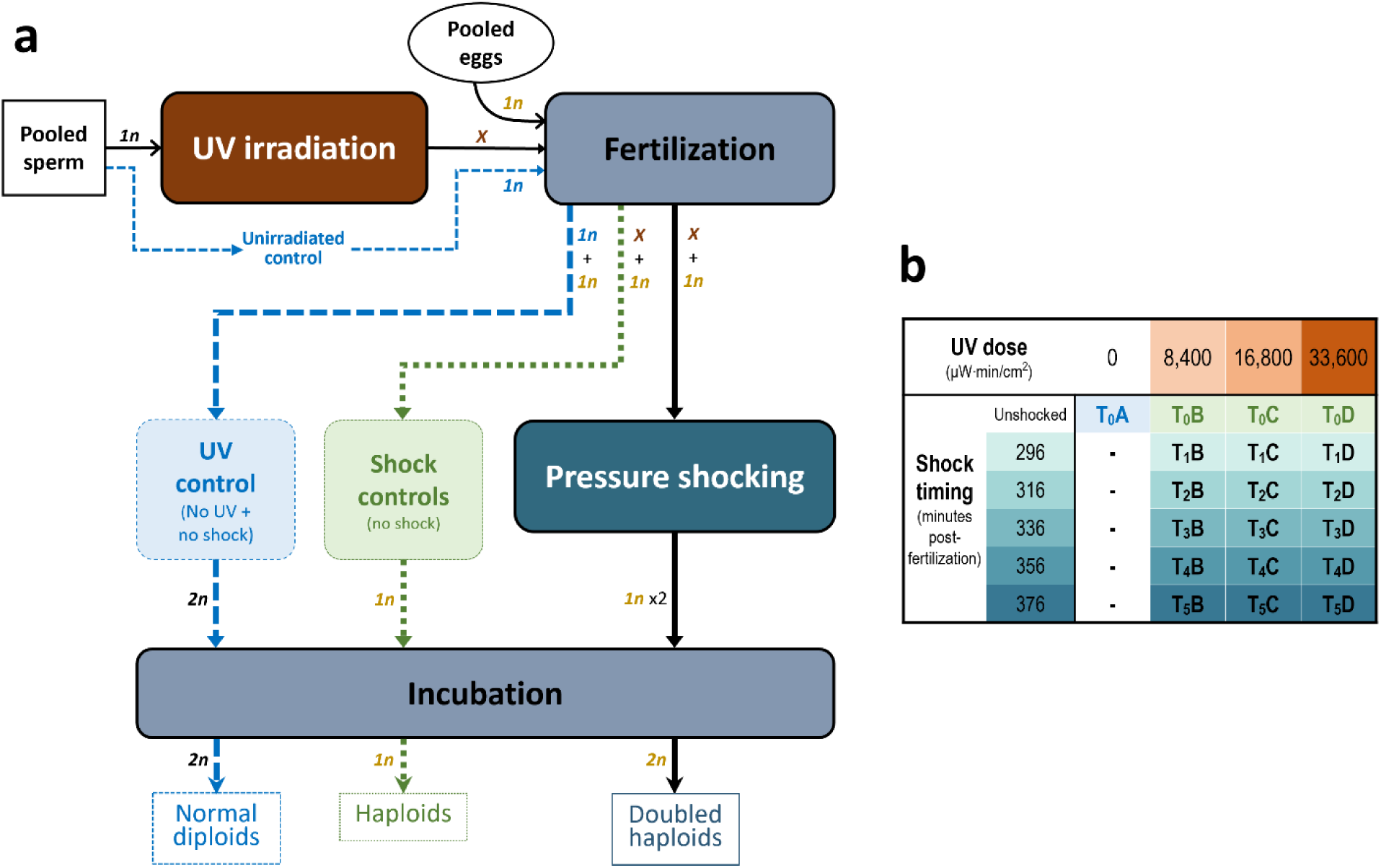
Overview of the mitotic gynogenic doubled haploid production procedure and of the experimental design. **a)** Sperm DNA was first inactivated through UV irradiation of milt. Following fertilization, eggs were hardened at 10°C, then exposed to a 10,000 psi pressure shock for five minutes in order to retain duplicated maternal DNA before the first cleavage interval (occurring at 4,789 degrees minutes at 10°C; Weber et al., 2014) and thus achieve diploidy. Expected ploidy at a given step is indicated by *1n* (haploid), *2n* (diploid) or *X* (deactivated sperm DNA) tags. **b)** Three UV irradiation doses (8,400, 16,800 and 33,600 µW/cm^2^·min; groups B-C-D) and five pressure shock times (from 296 to 376 minutes after fertilization; groups T_0_-T_1_-T_2_-T_3_-T_4_-T_5_) were trialled, yielding 15 treatment groups and four control groups.

Eggs and milt were collected from the F1 captive-reared progeny of wild origin broodstock that originated from Cape Race, Newfoundland, a region with numerous, isolated brook trout populations whose ecology and genetics have been extensively studied since the 1980s (Hutchings, 1993; Danzmann et al., 1998; Wood et al., 2014). The F1 fish were reared at Concordia University’s Applied Sciences Hub Aquatic Facility from the fertilized egg stage until adult maturation; their unfertilized gametes were shipped on ice to the LARSEM without supplemented oxygen. Given the small milt volume available for each male, all milt samples were pooled together to buffer against warming during subsequent manipulations at above zero temperature, which could be detrimental to the fertilization potential of sperm (Alavi & Cosson, 2005). Spermatozoa motility was checked by visual inspection of milt in a drop of 0.9% saline solution under a microscope to assess the quality of milt. Pooled milt was then divided into four UV treatment groups (A, B, C and D; Figure 1b), and 1 mL of milt from each group was diluted into 9 mL of AquaBoost Dilutor (Cryogenetics).

To initiate egg fertilization without paternal contribution, sperm DNA was inactivated by UV irradiation (Yamashita et al., 1990). We built a UV box in which four Philips TUV 30W 1SL/25 lights were suspended over a shaking platform in a closed bin installed in a refrigerated chamber at 4°C. A vacuum was connected to the box for providing continuous cold air entry and preventing warmup, which stabilized ambient temperature at 6°C during the irradiation process. Milt to be irradiated was placed in standard 100 mm plastic Petri dishes secured onto the shaking platform, 5.5 cm beneath UV lights. The UV intensity dispensed in this setup was measured at 2,200 µW/cm^2^ using a calibrated UV dose meter (Blak-Ray J-225 Short Wave UV Meter; San Gabriel, California). Based on previous successful mitogenetic double-haploid production trials with *Oncorhynchus* species, a total UV dose of 16,800 µW/cm^2^·min reliably achieves sperm DNA inactivation (i.e., 5,600 µW/cm^2^ for 3 minutes; Quillet, Garcia & Guyomard, 1991; Christensen et al., 2018a; Biagi et al., 2022). We thus used this value as a target for brook trout and calculated the required irradiation duration for providing 50, 100 and 150% of this target dose using our setup. Groups B, C and D milt preparations were thus irradiated during 229 (half of the target dose), 458 (target dose) and 687 seconds (150% of the target dose) respectively. Group A milt was not irradiated and served as control for UV inactivation of sperm.

As done for milt, all available eggs were pooled together prior to fertilization, then split into four groups (A-B-C-D; Figure 1b). Each egg group was then fertilized with one of the four milt preparations for activating the eggs, which were then hardened at 10°C. Eggs were then disinfected in a 10 mL/L solution of Ovadine for 10 minutes, and 50 mL of group A eggs were directly placed into incubation to be used as the control group T_0_A (unirradiated milt and unshocked; Figure 1b). To retain duplicated maternal DNA for producing doubled haploid progeny, the fertilized eggs were pressure-shocked to suppress the first mitotic division following fertilization. For *Oncorhynchus*, the optimal timing for performing pressure shocking is reportedly at around 60-70% of the first cleavage interval (FCI; time from fertilization to first mitotic division), which shows both inter and intraspecific variation (Hershberger & Hostetler, 2005). The three egg batches fertilized with UV-irradiated milt (groups B-C-D) were thus split into five groups (T_0_-T_1_-T_2_-T_3_-T_4_-T_5_) to test five shock times, i.e., at 70% of the FCI estimated for brook trout (4,789 °C·min at 10°C; Weber et al., 2014), and 20 and 40 minutes both before and after 70% of FCI time (Figure 1b). Each fertilized egg batch was shocked at 10,000 psi for 5 minutes at the determined shock time in a 2.7L-cylinder hydraulic pressure chamber (TRC Hydraulics Inc.), then stored in vertical incubators (MariSource 8-tray). Additionally, 50 mL of each UV-irradiated milt egg batch were not shocked to serve as controls.

Over the next few days, the incubation temperature was gradually reduced to 6°C and eggs were inspected daily for fungal development and mortality. Eggs showing signs of non-viability were discarded daily, whereas eggs exhibiting abnormal development features (e.g., apparent microphthalmic embryo) were pooled together separately from normal eggs. Alevins from each group were reared separately in controlled environment basins and were fed Skretting NutraXP (0.5, 0.7, 1.0 and 1.2 mm), Skretting Nutra RC (1.8, 2.3, 3.0 mm) and Bio-Oregon BioBrood. To prevent bacterial infection, chloramine T treatments (15 mg/L) were dispensed every 2-3 days over a month period following hatching (from April 26 to May 16). At eight weeks following hatching, 20 putatively doubled haploid alevins were collected for salt extraction of DNA (Aljanabi & Martinez, 1997) to test for homozygosity at 11 microsatellites using Qiagen’s Multiplex PCR kit (Supplementary Data 1 & 2; Dehaan & Ardren, 2005; Perry et al., 2005; King et al., 2012), which confirmed that all 20 alevins were indeed doubled haploids. Twenty months after hatching, heart, liver and muscle tissue were sampled from a doubled haploid female and stored at -80°C until DNA extraction.

### Sequencing

For long-read sequencing, high molecular weight DNA was extracted from 100 mg of heart tissue grinded in liquid nitrogen using Qiagen’s Genomics Tips 100G kit. Four libraries were prepared following the HiFi library preparation Express protocol, using 10 μg of (sheared) input DNA. Sequencing was then performed on a Pacific Biosciences Sequel II device at McGill University using eight SMRT 8M cells, with a target coverage of 120X. Primary raw data processing steps, including basecalling, adapter removal and circular consensus sequence (CCS) reads generation (ccs mode), were performed with SMRT Link 11.0.0.

To provide long-range information for guiding assembly, we also generated Hi-C data from the same individual sampled for PacBio sequencing. One proximity ligation library was prepared from 1,000 mg of grinded frozen muscle tissue using the Arima High Coverage HiC+ kit and the final Hi-C library was constructed using the NxSeq AmpFREE Low DNA Library kit. The Hi-C library was then sequenced on the MGI Technologies DNBSEQ-G400 using two lanes with PE150 cycles. Before sequencing, the library was first circularized using the MGIEasy Universal Library Conversion kit (App-A) followed by rolling circle amplification (RCA) to form the DNA nanoballs. The DNA nanoballs were then loaded in the flowcell before sequencing.

We also produced high coverage (70-80X) short-read data to be used for polishing the assembly. An additional high-molecular weight DNA extraction was performed using Qiagen’s Genomics Tips 100G protocol, with 80 mg of liver tissue grinded in liquid nitrogen as the input sample. Two PCR-free shotgun libraries were prepared from the DNA sample, which were then loaded on 0.326 of an S4 PE150 lane for sequencing on an Illumina NovaSeq6000.

All sequencing was performed at the McGill Genome Centre.

### Genome assembly

Genome assembly was conducted in four main steps, summarized in Figure 2. The different assembly versions generated throughout the procedure were compared to each other using QUAST version 5.0.2 (Gurevich et al., 2013) with default parameters. First, we produced a draft assembly from the raw PacBio ccs files using Flye with default parameters (version 2.8), a graph-based and repeat-aware assembler (Kolmogorov et al., 2019).

**Figure 2.**
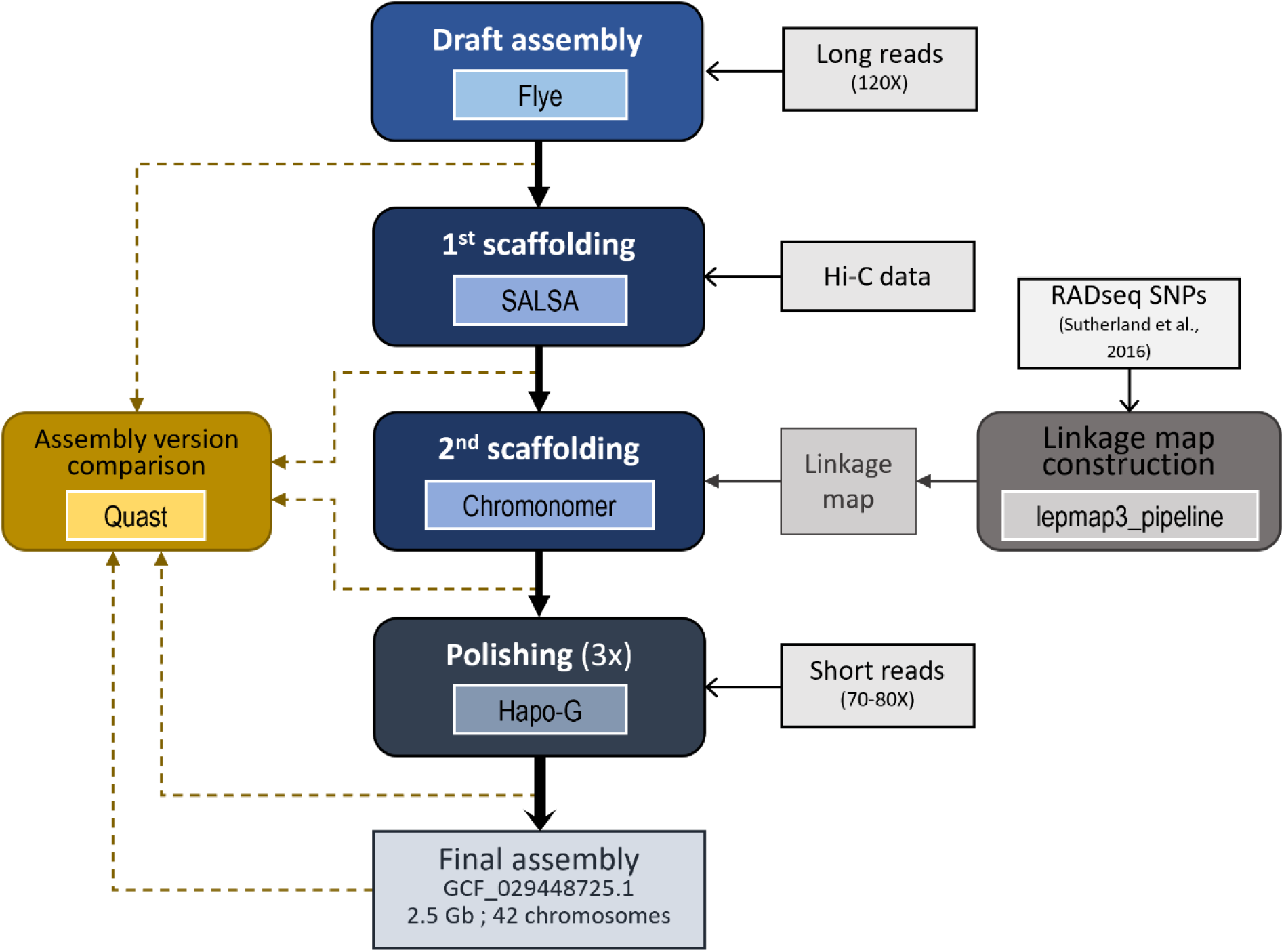
Overview of the brook trout genome assembly procedure and inputs. Rounded corner rectangles indicate processing steps, where inner boxes show the tools used for a given step. Inputs are shown in rectangles connected to other boxes with open arrows.

To scaffold the draft assembly, we mapped Hi-C reads to the draft assembly using the Arima Genomics Mapping pipeline (Arima Genomics https://github.com/ArimaGenomics/mapping_pipeline), then ran SALSA version 2 (Ghurye et al., 2017) with default parameters and mapped reads. We then performed an additional scaffolding procedure to anchor contigs into chromosomes by building a linkage map from previously published data (Sauvage et al. 2012a, 2012b, Sutherland et al., 2016) using Lep-MAP3 (Rastas, 2017). This map is based on a F2 family derived from an F0 captive-bred female from a wild anadromous population (Laval River, Forestville, Québec), and from an F0 male from a local domestic aquaculture strain bred at the Pisciculture de la Jacques-Cartier (Cap-Santé, Québec) (see Sauvage et al., 2012a and 2012b for experimental details). The 192 F2 full sibs and their two parents (F1) were sequenced with reduced representation sequencing (RADseq) by Sutherland et al. (2016). Raw reads were aligned on the draft contigs outputted by Flye using BWA MEM with default parameters (Li & Durbin, 2009). Following the guidelines provided in the Lep-MAP3 documentation (Rastas, 2017), genotype likelihoods were obtained with SAMtools *mpileup* (Danecek et al., 2021), keeping only positions with at least 3X coverage. A linkage map was then built using a custom Lep-MAP3 pipeline available at https://github.com/clairemerot/lepmap3_pipeline. First, markers exhibiting extreme segregation distortion (*χ^2^* test, *P* < 10^-12^), with more than 30% of missing data or that were uninformative (i.e., homozygous) were excluded with the *Filtering2* module. Remaining markers were assigned to linkage groups (LGs) using the *SeparateChromosomes* module with a logarithm of odds (LOD) of 26 and a minimum size of 100 markers. Markers unassigned to LGs were subsequently joined to LGs using the module *JoinSingle* with decreasing values of LOD until LOD=15 and a minimum LOD difference of 5. This procedure assigned 18,548 markers into 42 LGs. Within each LG, markers were ordered with 6 iterations of the *OrderMarker* module. The marker order from the run with the best likelihood was retained, then refined 6 times with the *evaluateOrder* flag with 5 iterations each. When more than 50 markers were plateauing at the same position, all of them were removed, as suggested by Lep-MAP3 guidelines (https://sourceforge.net/p/lep-map3/wiki/LM3%20Home/#ordermarkers2). Exploration for more stringent filtering or different values of LOD resulted in highly similar collinear maps. This procedure yielded a linkage map collinear to the one previously produced by Sutherland et al. using JoinMap4 (van Ooijen, 2006), but based on a higher density of markers (16,203; see comparison in Supplementary Fig. 1 done with MapComp (Sutherland et al., 2016)). Scaffolds were then assembled into chromosomes using Chromonomer with default parameters (version 1.13; Catchen, Amores, & Bassham, 2020), which anchors and orients scaffolds based on the order of markers in the linkage map.

The dual-scaffolded assembly was then polished using high-coverage Illumina short reads. Raw reads were first processed with fastp version 0.20.0 (Chen et al., 2018) to remove adapters, trim one base at each read tail (--trim_tail1=1 and trim_tail2=1) and to perform base correction in overlapped regions (--correction), as well as to filter out reads shorter than 120 bp (--length_required=120) or with a quality phred score lower than 28 (--qualified_quality_phred=28). The trimmed and filtered short reads were then supplied to Hapo-G version 1.3.3, which leverages haplotype and sequence information to correct assembly (Aury & Istace, 2021). This polishing procedure was repeated three times, each time using the newest version of the polished assembly produced at the previous Hapo-G run.

Following the NCBI submission validation procedure, bacterial contamination sequences were removed from a few contigs and scaffolds smaller than 10 kb were discarded. Assembly statistics were computed using assembly stats 1.0.1 (https://github.com/sanger-pathogens/assembly-stats) and BBTools 36.92 (Bushnell, 2014).

### Functional annotation

Functional annotation of the final assembly was accomplished through the NCBI Eukaryotic Genome Annotation Pipeline version 10.1 upon submission. Briefly, transcripts from various sources (RefSeq, GenBank, SRA and proteins) for brook trout and relevant species were mapped with BLAST to the assembly, in which repeats were previously masked with Windowmasker (Morgulis et al., 2006). Homology-based and *ab initio* gene prediction models were then performed from transcripts alignments using the NCBI’s Gnomon procedure (Souvorov et al., 2010), which produced a list of genes, mRNA, CDS and ncRNA. Genome completeness was then evaluated from the annotated gene set (primary assembly) using BUSCO v4.1.4 (Manni et al., 2021) in “protein” mode with the actinopterygii_odb10 lineage dataset.

### Repeated content identification

To further characterize repeated regions and transposable elements of the assembly, we first identified repeat contents using RepeatModeler version 2.0.1 (Flynn et al., 2020). We then extracted repeat entries for the Salmonidae family from RepeatMasker’s repeat database (Dfam_Consensus-20181026, RepBase-20181026) using the queryRepeatDatabase.pl utility, and merged these libraries with the custom library. This expanded custom library was then supplied to RepeatMasker (version 4.0.8; Smit, Hubley & Green, 2013) to hard-mask repeats in the assembly and classify repeated contents.

### Identification of homeolog chromosomes and homology relationships with other salmonid species

Salmonid genomes feature homeolog chromosomes due to the past whole-genome duplication event (Allendorf & Thorgaard, 1984; Crête-Lafrenière et al., 2012; Lien et al., 2016). To identify these homeolog regions, we mapped each chromosome to itself and all other chromosomes using *nucmer* (built-in mapper in MUMmer version 4.0.0; Marçais et al., 2018) with the --maxmatch parameter. For this purpose, the assembly was hard masked for repeats and scaffolds unanchored into chromosomes were removed. The alignment output was then formatted with the *show-coords -dlTH* command, and supplied to SyMAP (Soderlund, Bomhoff & Nelson, 2011; version 4.2 build 120) to detect collinear blocks using parameters min_dots = 30, top_n = 2 and merge_blocks = 1. The resulting collinear block coordinates were plotted using the *circlize* package (version 0.4.16; Gu et al., 2014) in R version 4.3.4 (R Core Team, 2024). Overlapping blocks of coordinates in each chromosome were merged with bedtools *merge* (Quinlan & Hall, 2010) to estimate the number of base pairs covered by at least one collinear block in each chromosome. These self-synteny blocks were remapped to the masked genome with LASTZ (version 1.04.15; Harris, 2007) to estimate sequence similarity by 1-Mb windows.

Given the whole genome duplication, most regions were expected to have at least one homolog region or chromosome in the genome. Consequently, regions that both lacked such a corresponding chromosome and exhibited elevated read coverage likely represented duplicated sequences collapsed during the assembly procedure. To characterize putatively collapsed regions, we mapped the short reads previously used for assembly polishing to the hard-masked chromosomes with bwa-mem and computed mean read depth by 1-Mb windows along the genome with mosdepth (Pedersen & Quinlan, 2017). As described in Mérot et al. (2023), regions with a read depth higher than the sum of the mean genome-wide read depth plus twice the standard deviation and that had no homologous chromosome were treated as putatively collapsed regions.

Besides self-synteny, we characterized homologous chromosomes and chromosomal rearrangements relative to sister species lake trout (*Salvelinus namaycush*; SaNama_1.0; GCA_016432855.1; Smith et al., 2022), as well as to Atlantic salmon (*Salmo salar*; ICSASG_v2; GCA_000233375.4; Lien et al., 2016). The ICSASG_v2 version of the Atlantic salmon genome was chosen to facilitate comparison with the chromosome correspondence previously established by Sutherland et al. (2016). Both assemblies were hard masked with RepeatMasker, using the custom library used for repeat-masking the brook trout assembly. Collinearity between the chromosomes of brook trout and those of the two species chosen for comparison was inferred using ntSynt (Coombe et al., 2024) with parameters -d 10 --block_size 1000000 --indel 100000 --merge 1000000 --w_rounds 500 250. Interspecific syntenic relationships were then plotted with a custom version of the *draw.linear()* function of the *syntenyPlotteR package* (version 1.1.0; Quigley et al., 2023).

## Results and discussion

### Doubled haploid production

Despite low survival, some combinations of treatments successfully resulted in viable doubled haploids. At eight weeks post-fertilization, mortality was moderately high (79.15%; Supplementary Table 1) in the control group T_0_A (unirradiated milt and unshocked), which could be attributed to the various factors, including transportation conditions and the fact that eggs were derived from the F1 captive-reared progeny of a wild-origin broodstock known to exhibit poor performance in hatchery conditions (Fraser et al., 2019). Mortality ranged from 83 to 95% in the egg groups that were both fertilized with UV-irradiated milt and pressure-shocked, hence putatively doubled haploids. Mortality was noticeably high (90 to 97%) among putatively haploid egg groups, i.e., unshocked eggs fertilized with UV-irradiated milt; groups T_0_B-C-D). Similarly, approximately half of eggs from each of these unshocked and UV-treated milt groups bore visibly abnormal microphthalmic (“pin-eyed”) embryos; such abnormalities in eye development are often found in haploid individuals (R. Devlin, personnel communication; Arai, 2001; Luo & Li, 2003; Arai & Fujimoto, 2018). Conversely, the proportion of pin-eyed eggs among the putatively double haploid groups (UV-treated milt and shocked eggs) was at most 25% (T_1_D). These observations strongly suggest that UV irradiation was successful in inactivating sperm DNA and resulted in haploid embryos in unshocked egg groups, which rarely survive until hatching.

During the period between weeks 8 and 15 after fertilization, remaining viable eggs exhibiting normal development from all three UV-treated milt and shocked groups (B-C-D) were pooled together by shock time (T_1_ to T_5_). At the 15^th^ week post-fertilization, mortality was the lowest (1.61%) in the untreated control group T_0_A, and considerably higher in putatively haploid egg groups (unshocked and UV-irradiated milt egg groups T_0_B, C and D) (82-97%; Supplementary Table 2), whereas mortality in putatively doubled haploid egg groups ranged from 25% (T_5_B-C-D) to 35% (T_1_B-C-D). These candidate doubled haploid alevins, along with normal diploid control alevins (T_0_A), began hatching about 70 days after fertilization. Some of the putative doubled haploid alevins had enlarged yolk sacs, a known developmental abnormality of doubled-haploid salmonids (Jagiełło et al., 2017; Jagiełło et al., 2021). None of the 205 pin-eyed eggs from all treatment groups (which were pooled together into a single group, B-C-D pin-eyed; Supplementary Table 2) survived until hatching, suggesting that these pin-eyed eggs were indeed haploid and therefore that either sperm DNA inactivation and/or that retention of duplicated maternal DNA failed in a few eggs of each group.

Of the 773 candidate doubled haploid alevins that hatched, 49% survived to week 25 post-fertilization. Those candidates represented 88% of all the surviving larvae across all treatment and control groups. Many featured various morphological deformities described by Jagiełło et al. (2017; 2021), mostly cephalic and spinal defects. Subsequent genotyping of 20 of these individuals at 11 microsatellite loci confirmed they were indeed all doubled haploids.

Because viable potential doubled haploid alevins from different treatment groups were pooled together post-hatching, it is not possible to investigate the effect of different UV doses or shock times on doubled haploid alevin viability, which would have required considerably more space and was not the aim of the current study. This would also have required consistent UV intensity and temperature readings, which showed important variations in our irradiation setup. However, the experimental parameters used here may serve as guidelines for future doubled haploid production in salmonids. We hypothesize that cooling milt during UV irradiation was critical for preserving sperm’s ability to fertilize eggs while successfully inactivating sperm DNA and very likely contributed to successful mitogynogenesis.

### Genome assembly

We generated a reference genome assembly for brook trout through multiple rounds of scaffolding and refinement using various tools and sequencing data inputs.

The initial draft assembly produced from high coverage PacBio reads had a size of 2.498 Gb and featured 9,083 scaffolds (Table 1). Scaffolding the draft assembly with Hi-C data reduced the number of scaffolds by almost half to 4,737 and increased N50 by almost seven times to 8,184,474 bp. The incorporation of linkage map information at the second scaffolding step further refined the assembly into 4,062 scaffolds, including 42 linkage groups, which accounted for 88.67% of the assembly’s total size (2.502 Gb). While the assembly obtained at this stage had the largest size and N50 among all five versions summarized in Table 1, it also had the most gaps (5,227).

**Table 1.** Assembly statistics for versions obtained at different stages of the brook trout genome assembly procedure. Chromosome scale corresponds to the proportion of the total assembly length that is anchored into main chromosomes. NA indicates a value that could not be estimated at the given stage (i.e., prior to scaffolding into chromosomes or determination of gaps and N nucleotides).

|  | Assembly stage |  |  |  |  |
| --- | --- | --- | --- | --- | --- |
|  | Draft | 1<sup>st</sup> scaffolding | 2<sup>nd</sup> scaffolding | Polished | Final |
| **Total length (bp)** | 2,497,860,955 | 2,502,111,779 | 2,502,179,279 | 2,501,987,914 | 2,496,201,733 |
| **Total number of scaffolds** | 9,083 | 4,737 | 4,062 | 4,062 | 2,721 |
| **N50** | 1,190,200 | 8,184,474 | 50,987,102 | 50,983,633 | 50,983,633 |
| **L50** | 537 | 64 | 19 | 19 | 19 |
| **N90** | 158,250 | 458,326 | 876,059 | 875,901 | 985,419 |
| **L90** | 2,730 | 583 | 66 | 66 | 60 |
| **Chromosome scale** | NA | NA | 0.887 | 0.887 | 0.889 |
| **Longest scaffold length (bp)** | 7,916,649 | 53,452,995 | 101,322,550 | 101,316,066 | 101,316,066 |
| **Smallest scaffold length (bp)** | 23 | 23 | 23 | 23 | 10,004 |
| **Missingness (N %)** | NA | 0.002 | 0.002 | 0.002 | 0.002 |
| **Gaps** | NA | 4,603 | 5,227 | 4,309 | 4,299 |
| **GC content** | 0.437 | 0.437 | 0.437 | 0.437 | 0.437 |

The three rounds of polishing using Illumina short reads resulted in fewer gaps (4,309), but also in a slightly smaller assembly size and N50. However, given the higher base accuracy of short reads compared to long reads (Delahaye et al., 2021), polishing with short reads likely improved sequence consensus compared to the scaffolded version, which is desirable for reliable gene prediction (Watson & Warr, 2019; Aury & Istace, 2021).

A total of 1,341 scaffolds with lengths under 10 kb were excluded through the NCBI submission process.The final assembly comprises 42 primary chromosomes, which represent 88.88% of total assembly size (2.496 Gb), and 2,679 unplaced scaffolds, including the mitochondria (scaffold CM055727.1, 16,623 bp), with an N50 of 50,983,633. The proportion of N nucleotides and GC contents remained the same among all assembly versions.

The final assembly length (2.496 Gb) falls within the 2-3 Gb size range typically expected for salmonid genomes (Supplementary Table 3). It is smaller than the assembled genomes of Atlantic salmon (*Salmo salar*; 2.967 Gb) and lake whitefish (*Coregonus clupeaformis*; 2.753 Gb) but larger than the rainbow trout (*O. mykiss*; 2.342 Gb) and river trout (Salmo trutta; 2.372 Gb) assemblies. The brook trout assembly is also the largest among the most closely related charr species such as lake trout (*S. namaycush*; 2.345 Gb) and Dolly Varden (*Salvelinus* sp.; 2.170 Gb), of which the former was assembled using a hybrid approach similar to the assembly procedure conducted for the brook trout genome (e.g., Hi-C, long- and short-read sequencing) (Smith et al., 2021). While part of this variation in salmonid genome size might be due to technical artifacts during assembly, fish genomes are known for their genome size heterogeneity, linked to polyploidization and rediploidization events, but also to ecological characteristics (Hardie & Hebert, 2004). In particular, it is worth noting that larger genomes have been associated with greater ecological tolerance in fish (Hardie & Hebert, 2004), and besides having the largest genome among chars, brook trout is known to display considerable plasticity in life history characteristics, such as body size and morphology (Thériault et al., 2007; Morinville & Rasmussen, 2008).

### Genome annotation

Through the NCBI Eukaryotic Annotation Pipeline, 56,058 genes (Table 2) were identified, with an average length of 23.67 kb, as well as 83,302 CDS, 487,554 exons and 487,554 introns. These genomic features were found in greater numbers in the brook trout assembly than in related charr assemblies (Supplementary Table 4). In addition, 99.0% of the 3,640 expected conserved orthologs BUSCO genes were found complete within the genome annotation of the brook trout assembly, which strongly suggests an elevated assembly completeness, with only 1% of BUSCO genes fragmented or missing. The assembly also features one of the highest proportions of complete and single copy BUSCO genes (57.3%) reported across all six salmonid species selected for comparison (Supplementary Table 4).

**Table 2.** Functional annotation of genomic features of the brook trout genome assembly obtained from the NCBI Eukaryotic Annotation Pipeline. Total gene count excludes pseudogenes.

| Annotated features |  |  |
| --- | --- | --- |
| Number of genes | 56,058 |  |
| Average gene length (bp) | 23,670 |  |
| Number of protein-coding genes | 41,382 |  |
| Number of CDS | 83,302 |  |
| Average CDS length (bp) | 2,259 |  |
| Number of exons | 487,554 |  |
| Average exon length (bp) | 394 |  |
| Number of introns | 435,191 |  |
| Average intron length (bp) | 3,280 |  |
| BUSCO genes |  |  |
| Complete and single copy | 57.3 |  |
| Complete and duplicated | 41.7 |  |
| *Complete* |  | 99.0 |
| Fragmented | 0.5 |  |
| Missing | 0.5 |  |

### Repeated content identification

As expected for salmonid genomes, over half of genome base pairs were masked as repeats (1,583,475,168 bp; 63.44%). These repeats were mostly unclassified elements (23.00%; Table 3). 1,461,010 DNA transposons were identified (19.22%), mainly Tc1-IS630-Pogo, and most of the retroelements identified were LINEs (16.71%). Overall, interspersed repeats account for 58.93% of the genome (1,471,091,031 bp).

**Table 3.** Number of elements, total sequence length and percent of genome length representing repeated elements in the brook trout genome.

|  | Number of<br>elements | Total length<br>(bp) | Fraction<br>of total<br>length (%) |
| --- | --- | --- | --- |
| **Retroelements** | 1,134,327 | 417,075,081 | 16.71 |
| SINEs | 194,163 | 33,890,667 | 1.36 |
| Penelope | 24,565 | 5,105,541 | 0.20 |
| LINEs | 744,848 | 292,805,214 | 11.73 |
| L2/CR1/Rex | 525,213 | 213,268,889 | 8.54 |
| R1/LOA/Jockey | 14,915 | 3,955,522 | 0.16 |
| R2/R4/NeSL | 3,353 | 950,188 | 0.04 |
| RTE/Bov-B | 81,441 | 32,408,147 | 1.30 |
| L1/CIN4 | 25,777 | 12,649,380 | 0.51 |
| LTR elements | 195,316 | 90,379,200 | 3.62 |
| BEL/Pao | 11,947 | 2,817,887 | 0.11 |
| Ty1/Copia | 34,864 | 15,420,579 | 0.62 |
| Gypsy/DIRS1 | 103,202 | 49,625,102 | 1.99 |
| Retroviral | 31,402 | 11,789,266 | 0.47 |
| **DNA transposons** | 1,461,010 | 479,842,353 | 19.22 |
| hobo-Activator | 325,779 | 81,181,519 | 3.25 |
| Tc1-IS630-Pogo | 581,758 | 244,783,508 | 9.81 |
| PiggyBac | 20,749 | 6,910,307 | 0.28 |
| Tourist/Harbinger | 33,826 | 7,552,238 | 0.30 |
| Other (Mirage, P-element,<br>Transib) | 2,337 | 232,926 | 0.01 |
| **Rolling-circles** | 24,196 | 7,495,875 | 0.30 |
| **Unclassified** | 1,806,852 | 574,173,597 | 23.00 |
| **Total interspersed repeats** |  | 1,471,091,031 | 58.93 |
| **Small RNA** | 12,570 | 3,920,647 | 0.16 |
| **Satellites** | 62,235 | 46,250,500 | 1.85 |
| **Simple repeats** | 517,969 | 50,835,562 | 2.04 |
| **Low complexity** | 62,311 | 6,476,267 | 0.26 |

### Identification of homeolog chromosomes and homology relationships with other salmonid species

Self-alignment of brook trout chromosomes revealed 384 synteny blocks of varying sizes, ranging from 200 kb to 40.6 Mb (Figure 3). These blocks accounted for 97.8% of genome base pairs anchored into the 42 main chromosomes, and all chromosomes had at least one homeolog region. However, for some smaller chromosomes (34, 35, 39, 40 and 42), the proportion of base pairs matched to a homeolog region was under 75% (Supplementary Table 5), especially in telomeric regions. Additionally, we identified only fourteen putatively collapsed regions (10.078 Mb; Supplementary Table 6) for which no homeologs were found and that exhibited increased read coverage, fewer than reported for Atlantic salmon (Lien et al., 2016) and for dwarf lake whitefish (Mérot et al., 2023), which likely results from the use of a doubled haploid sample. These putatively collapsed regions mostly appear to be in or near telomeres, which is expected since telomeric regions are known to show residual tetraploidy (Allendorf et al., 2015), leading assembling tools to merge those similar sequences in a single reference sequence.

**Figure 3.**
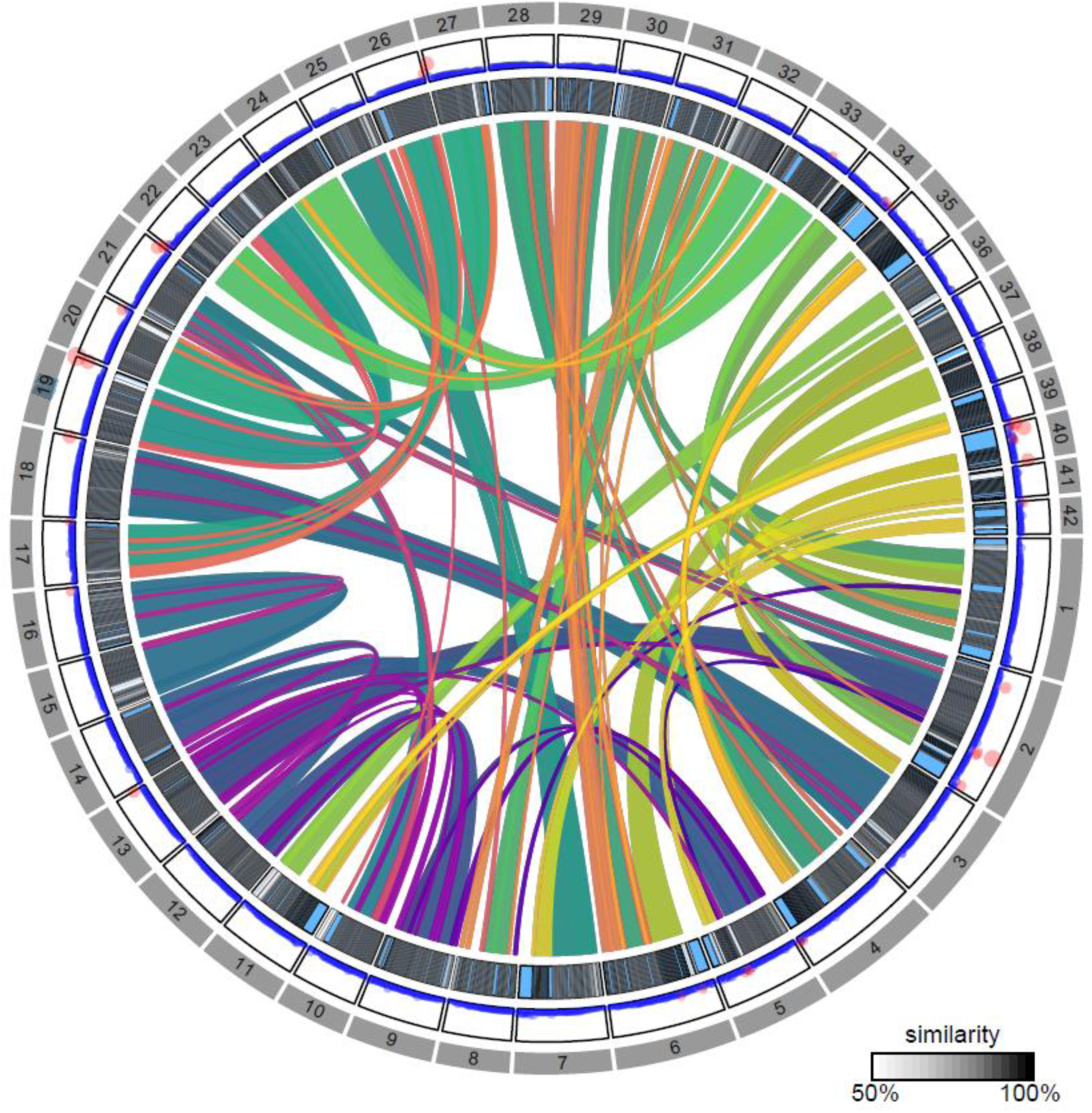
Self-synteny, similarity of homeolog regions and read coverage along the 42 chromosomes of the brook trout (*S. fontinalis*) genome. The inner track represents relationships between homeologous chromosomes and the middle track shows their degree of similarity, where pale blue areas are regions for which no homology was found. The outer track displays short-read coverage by 1-Mb windows. Red dots indicate windows where read coverage exceeds the sum of the mean genome-wide read depth and twice the standard deviation. Areas with elevated coverage and no homology are putatively collapsed regions.

When comparing with other species, chromosomes appeared highly conserved between lake and brook trout, as all of their 42 chromosomes were at least partially collinear with each other (Figure 4). While collinearity was restricted to some regions for some small chromosomes (34, 40), most brook trout chromosomes were collinear with a given lake trout chromosome across almost all their length. This strong collinearity between both *Salvelinus* species is consistent with them belonging to the same salmonid lineage. Conversely, homolog relationship between brook trout and Atlantic salmon genomes were characterized by multiple chromosomal rearrangements relative to the brook trout genome, which is expected given the greater phylogenetic distance between the two species and their different number of chromosomes. Indeed, for many salmon chromosomes, we found collinear regions distributed among two or three distinct brook trout chromosomes corresponding to the species-specific fusions and fissions previously inferred from linkage maps by Sutherland et al. (2016). For example, chromosomes 1 and 9 of salmon each comprise three arms (Philips et al., 2009) that appeared to be derived from tandem fusions of three brook trout chromosomes, e.g., 1 (BC1), 15 (BC12), 21 (BC13), and 16 (BC35), 29 (BC33), 33 (BC38), respectively. While we found one-to-one homologous relationships with salmon chromosomes for brook trout chromosomes 12, 21, 22 and 28, most of salmon chromosomes apparently result from the fusion of two ancestral post-whole-genome duplication chromosomes that remain distinct linkage groups in brook trout. However, for some regions and chromosome arms, we found little to no correspondence with any region of the other species’ genome, such as brook trout chromosome 34 and one arm of salmon chromosome 3, which might indicate more interspecific divergence in those regions. These regions may thus represent candidates of interest for brook trout-specific innovations and life history traits.

**Figure 4.**
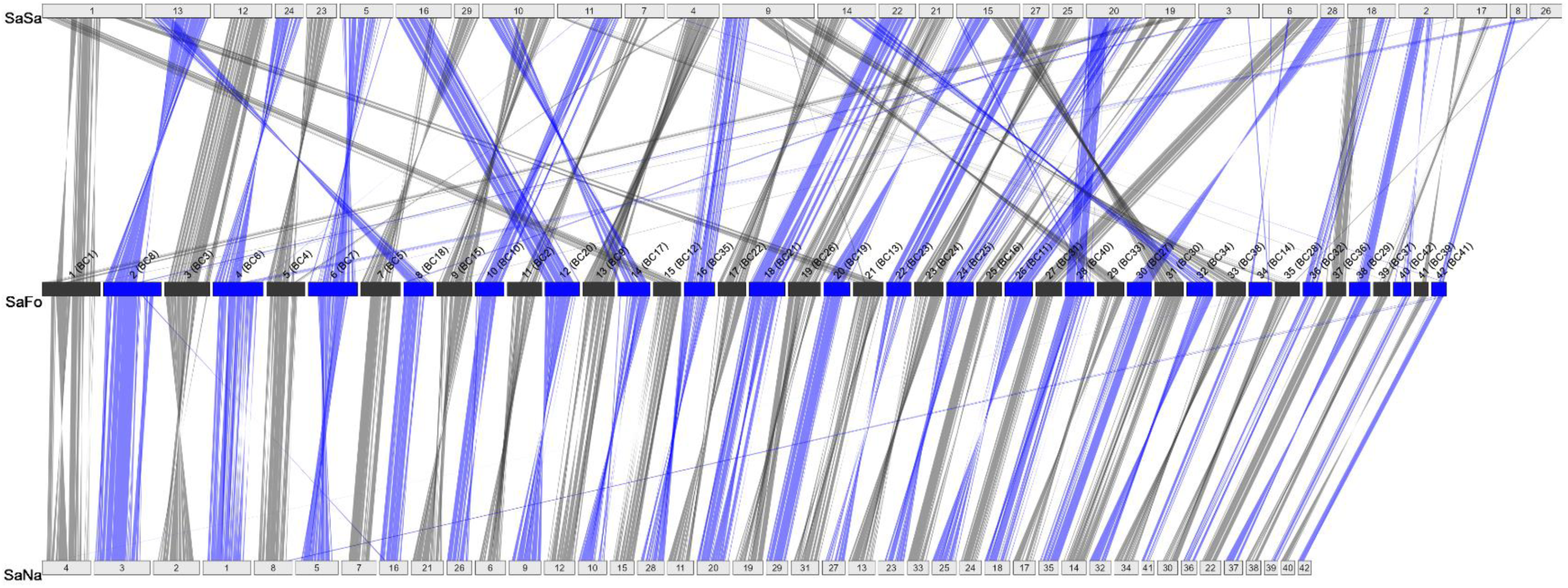
Syntenic relationships between chromosomes of brook trout (*Salvelinus fontinalis*; SaFo), lake trout (*Salvelinus namaycush*; SaNa) and Atlantic salmon (*Salmo salar*; SaSa). Parentheses next to brook trout chromosome numbers indicate corresponding linkage groups determined previously in Sutherland et al. (2016) (see Supplementary Table 7 for correspondence between linkage groups and chromosome numbers).

## Conclusion

We present a high-quality chromosome-level assembly for brook trout produced from PacBio CCS long reads, Illumina short reads and Hi-C sequencing data. Its elevated contiguity and completeness are attributable to the use of fully homozygous doubled haploid fish produced by mitotic gynogenesis. The assembly features 42 chromosomes that account for 88.88% of the 2.496 Gb assembly size and is comparable to other salmonid genome assemblies concerning quality metrics and the number of annotated genes features. Synteny analysis suggested minimal assembly collapse and confirmed previously established homologous relationship between the chromosomes of brook trout and Atlantic salmon. This assembly thus represents a novel tool that will allow exploration of various research avenues, including the role of transposable elements and repeats in genome evolution and adaptation, functional genomics and comparative genomics of salmonid chromosomes. It will also enable the use of genomic markers such as single nucleotide and structural variants to support sustainable management of harvested populations and future conservation.

## Data availability

The final brook trout genome assembly ASM2944872v1 is available from GenBank (accession GCA_029448725.1) under BioProject PRJNA922922. The assembly and its annotation are also available from RefSeq (accession GCF_029448725.1).

Tools and parameters used in each assembly step are detailed throughout the manuscript.

Raw sequencing data (long and short reads, Hi-C) was deposited in NCBI’s Sequence Read Archive (SRA) under BioProject PRJNA922922 and BioSample SAMN32676627.

Post-assembly analysis scripts can be found in the GitHub repository https://github.com/LaurieLecomte/safo_paper.

## Acknowledgments

We are thankful to Concordia University personnel for overseeing fish rearing, egg and milt collection and transportation, as well as to all members of the Laboratoire de recherche en sciences environnementales et médicales (LARSEM) at Université Laval who contributed to doubled haploid production. We are also very grateful to Robert Devlin for providing highly relevant information and lending equipment to assist with doubled haploid production, and Gabriel Piette-Lauzière and Maeva Leitwein for assistance with sampling and Cecilia Hernandez for the genotyping of doubled haploids. This project was initiated by Louis Bernatchez, who passed away during the production of this publication. We dedicate this article to his memory.

